# Impaired online error-correction disrupts synchronization to external events in autism

**DOI:** 10.1101/2020.09.28.316828

**Authors:** Gal Vishne, Nori Jacoby, Tamar Malinovitch, Tamir Epstein, Or Frenkel, Merav Ahissar

## Abstract

Autism is a developmental disorder characterized by impaired social skills and accompanied by motor and perceptual atypicalities. Its etiology is an open question, partly due to the diverse range of associated difficulties. Based on recent observations that individuals with autism are slow in updating perceptual priors, we now hypothesized that motor updating is also slow. Slow motor updating is expected to hamper the ability to synchronize to external events, since asynchronies are corrected sluggishly. Since sensorimotor synchronization is important for social bonding and cooperation, its impairment is expected to impair social skills. To test this hypothesis, we measured paced finger tapping to a metronome in neurotypical, ASD, and dyslexia groups. Dyslexia was assessed as a control group with a non-social neurodevelopmental atypicality. Only the ASD group showed reduced sensorimotor synchronization. Trial-by-trial computational modelling revealed that their ability to form controlled motor responses and to maintain reliable temporal representations was adequate. Only their rate of error-correction was slow and was correlated with the severity of their social difficulties. Taken together, these findings suggest that slow updating in autism contributes to both sloppy sensorimotor performance and difficulties in forming social bonds.

**Significance:** The prevalence of autism diagnosis has increased immensely is the last decades. Yet its etiology remains a challenge, partly since the functional relations between characteristic social difficulties, perceptual and motor atypicalities are not understood. Using trial-by-trial computational modelling, we show that a single deficit underlies the poor synchronization of individuals with autism in both static and changing environments. Slow updating, leading to slow online error correction of motor plans, has an immense explanatory power explaining both difficulties in sensorimotor synchronization, and social impairments.

## Introduction

The core difficulty in social interactions of individuals with ASD has traditionally been attributed to a lack of social interest and motivation (Chevallier et al., 2012), but this view has been recently challenged (Jaswal & Akhtar, 2019). Recent studies revealed that atypical perceptual and motor processing are consistent characteristics of autistic experience (Robertson & Baron-Cohen, 2017). For example, individuals with ASD show poor sensorimotor integration (Whyatt & Craig, 2013; Hannant et al., 2016a), whose magnitude is correlated with symptom severity (Hannant et al., 2016b). In parallel, there has been an emergent understanding that the act of synchronized activity contributes to social bonding, possibly by promoting a general sensorimotor predictive mechanism responsible for simulating others’ actions (Novembre et al., 2019). Thus, recent studies have found that moving in coordination with others increases pro-social helpful behavior (Kokal et al., 2011; Valdesolo & DeSteno, 2011; Tarr et al. 2014), enhances the ratings of affiliation (Hove & Risen, 2009), and strengthens the feelings of social bonding (Cirelli, 2018). The accumulative understanding of the importance of synchronous activity as an “engine” that promotes social communication led us to ask whether a single underlying mechanism could mediate both. We hypothesized that poor sensorimotor performance does not result from noisy motor characteristics, but rather from a more specific difficulty in integrating information online, which might reduce the rate of sensorimotor integration as well as impede some aspects of social cooperation. To test this hypothesis, we measured synchronization in the simple context of synchronized finger tapping, which also allowed us to assess and model synchronization in a non-social context.

Recent modelling of a perceptual task (Lieder et al., 2019), which characterized the dynamics of inference based on previous stimuli in the context of tone discrimination, found that individuals with ASD were markedly slower than neurotypical individuals in updating their perceptual predictions (priors). Based on this observation, we hypothesized that slow updating in individuals with ASD is a general characteristic of autism, which is also manifested in slower updating of motor plans. Accordingly, individuals with autism are predicted to have difficulties in synchronization to external stimuli due to slow online correction of their synchronization errors —their phase difference from the external rhythm. Thus, while slow updating predicts difficulties in sensorimotor synchronization, this is not due to “sloppy” motor commands or noisy retention of the underlying rhythm (Wing & Kristofferson, 1973). Rather, it suggests an impairment in the mechanism of online error-correction, which is necessary for fast corrections of small deviations from synchrony that otherwise accumulate to gradually increasing errors.

Paced finger tapping, in which the participant is asked to synchronize to the beat of an external metronome, offers two advantages for assessing the hypothesis of slower error correction. First, data acquisition is fast and reliable (test re-test correlation of the main tapping parameters is ∼.8; Figs. S1-S2). Second, it has been comprehensively modelled in the general population (Repp 2005; Repp & Su, 2013), though not in neurodevelopmental populations. In these models, the participant’s tapping is determined by motor noise, timekeeping noise, and error correction mechanisms (Wing & Kristofferson, 1973; Mates, 1994; Vorberg and Wing 1996; Vorberg and Schulze 2002; Jacoby et al., 2015). We administered two protocols using both a stationary environment (fixed metronome tempo, Experiment 1), and a volatile, changing environment (tempo-switch protocol, Experiment 2). In both experiments, perfectly synchronous behavior means perfect alignment between the participant’s taps and the external metronome beat. The error, defined as the temporal deviation between the participant’s response and the metronome beat, is termed asynchrony. Most individuals tap slightly ahead of each beat, though they hear their tap and the metronome as synchronous, hence the perceived synchrony is characterized by a small negative asynchrony (Aschersleben & Prinz, 1995; Repp 2005; Fig. 1a). The main challenge in this task is to keep the variability around this perceived synchrony very small. However, two sources of noise make the task difficult even when metronome tempo is fixed: noise in motor responses, and noise in the representation of the metronome tempo. Both can be corrected online by using the asynchrony signal (local deviations from the mean asynchrony). If errors are not corrected quickly and are kept through metronome beats, the variability around mean asynchrony accumulates to a large value and synchrony is poor. When the metronome tempo changes, the participant needs to correct for this change as well by quickly modifying the internal representation of the external tempo. We used this manipulation to test the complementary prediction of the “slow updating” hypothesis: reduced ability to quickly update to a changing environment. Importantly, the predictions of the slow-updating hypothesis contrast with predictions of other recent hypotheses, which propose that individuals with autism overestimate the volatility of the external environment (Lawson et al., 2017), and over-weigh their prediction errors (Van de Cruys et al., 2014). These hypotheses predict that external changes in the environment will be effectively corrected (even over-corrected) by individuals with autism, whereas the slow updating hypothesis postulates slower and locally reduced corrections.

**Figure 1:**
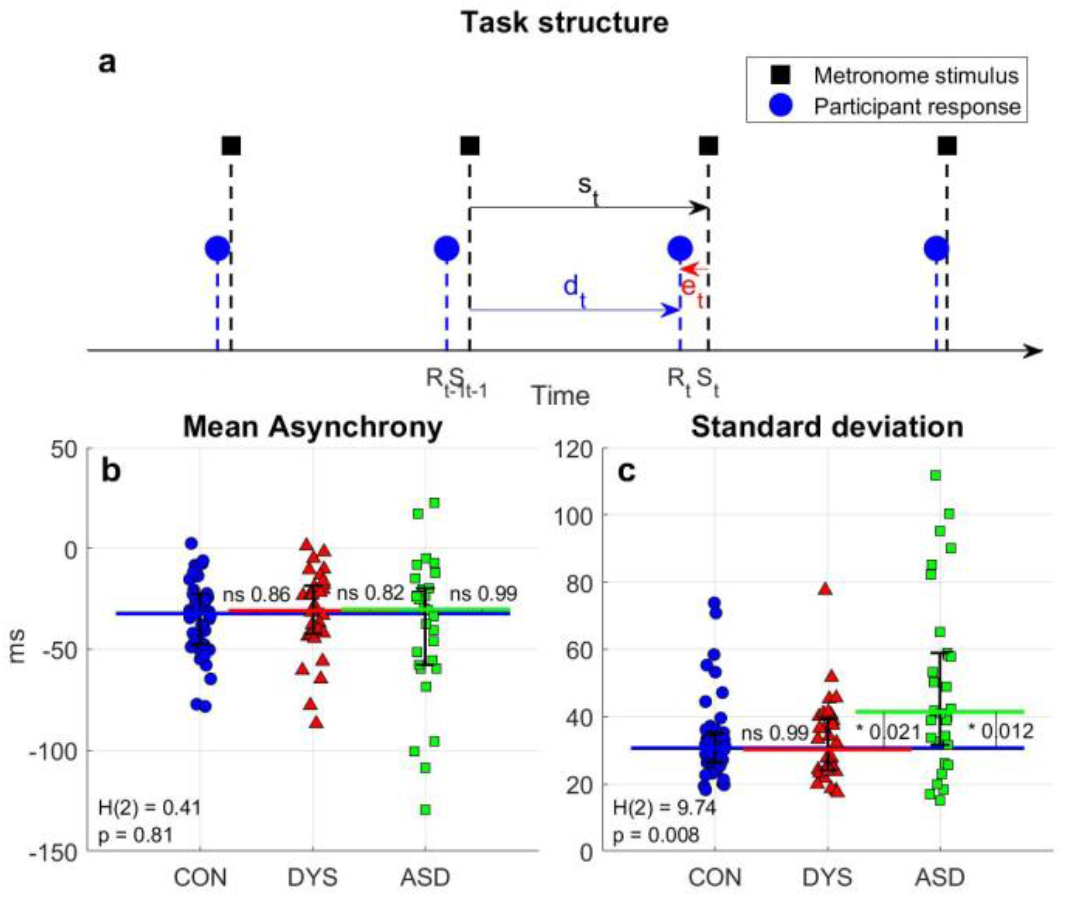
Isochronous finger tapping: mean asynchrony is similar in the three groups, but variability around this mean is substantially larger in ASD compared to neurotypical and dyslexia groups. (a) A schematic illustration of the temporal structure of paced tapping: metronome stimuli (black squares; *S*_*t*_ denotes t^th^ stimulus) and finger-tap responses (blue circles; *R*_*t*_ denotes t^th^ tap) as a function of time; *e*_*t*_ - error (asynchrony) in tap t; *s*_*t*_ - inter-beat interval; *d*_*t*_ - delay interval from the previous metronome stimulus to the following finger tap. (b-c) Basic tapping statistics: (b) Mean asynchrony is negative for all three groups (p < 10^*−*5^) and similar in the three populations, though more broadly distributed in the ASD group. (c) Standard deviation is larger in the ASD group compared to the two other groups. Each dot represents the performance of one participant (average of two blocks); y-axis represents the score in ms, and x-axis and color represent group membership (with a small jitter for readability): blue circles - neurotypical, red triangles - dyslexia, and green squares - ASD. The median of each group is denoted as a line of the same color; error bars denote interquartile range. Kruskal Wallis H-statistic and corresponding p-values are plotted in the bottom-left corner; p-values of comparisons between groups are plotted next to the line connecting the groups’ medians.

An important limitation when comparing individuals with autism to neurotypical individuals is setting criteria for a matched control group beyond age and general reasoning. Having a specific developmental difficulty is often accompanied with other atypical characteristics, like attentional deficits (e.g. high comorbidity of ASD and ADHD; Smith & Matson, 2010), and sometimes with associated emotional difficulties. Indeed, theories of underlying sensory and motor deficits have often been criticized for their lack of specificity (Goswami, 2015; Rogers & Ozonoff, 2005), i.e. for noting difficulties that characterize several neurodevelopmental disorders. To address this limitation, we recruited an additional control group from a population with a specific neurodevelopmental atypicality, namely individuals with developmental dyslexia. Individuals with dyslexia are characterized by poor reading and spelling in spite of intact intelligence and adequate learning opportunities (American Psychiatric Association, 2013). Like individuals with ASD, they show high concurrence with ADHD (Germano et al., 2010), and atypical perceptual characteristics (e.g. Ahissar et al., 2000; Tallal, 2004). A prominent hypothesis known as the “temporal sampling framework” posits that individuals with dyslexia have impaired oscillatory entrainment, which hampers linguistic processing (Goswami, 2011). This theory predicts that sensorimotor synchronization will also be impaired in dyslexia, since it also utilizes oscillatory neural mechanisms (Repp 2005; Repp & Su 2013). However, observations regarding synchronization in dyslexia are mixed (e.g. adequate tapping, Tiffin-Richards et al., 2004). Based on observations of adequate updating in dyslexia (Lieder et al., 2019), we hypothesized that fast online phase correction in dyslexia would be intact, and hence individuals with dyslexia will adequately synchronize to an external metronome. Experiments 1 and 2 tested these predictions.

## Results

### Experiment 1 – isochronous tapping reveals impaired online error correction in ASD, but not in dyslexia

We measured the mean asynchrony and variability in a paced finger-tapping task, with a fixed 2 Hz metronome beat (illustrated in Fig. 1a). We recruited three age and cognitive matched groups (Table S1) – neurotypical individuals (n=47), individuals with ASD (n=30) and individuals with dyslexia (n=32). As expected (Aschersleben & Prinz, 1995), the mean asynchrony manifested by most participants was negative (105/109 participants; 96.3%). Mean asynchrony was similar in the three groups (median [interquartile range] (ms): neurotypical: −32.2 [24.9], dyslexia: −30.8 [24], autism: −30.3 [37.9], Kruskal Wallis test H(2)=0.41, p>0.8, Fig. 1b), suggesting that the overall perceptual accuracy of the temporal intervals is similar in the three groups.

By contrast, we found a large difference in the variability (denoted by the standard deviation - SD) of the groups around their mean asynchrony (median [interquartile range] (ms): neurotypical: 30.6 [8.9], dyslexia: 30.2 [15.6], autism: 41.4 [27.1], Kruskal Wallis test H(2)=9.74, p=0.008, see Fig. 1c). The significant group difference was due to the large variability of individuals with ASD (post-hoc analysis of ASD group vs. both other groups using Tukey-Kramer method (throughout the paper): p<0.022, Cliff’s delta > 0.38 in both cases), while there was no difference between the dyslexia group and the neurotypical group (p>0.95). Although there were individuals with autism whose SD was in the range of the neurotypical population, the SD of a third of the group was more than two SDs above the neurotypical mean, compared with only one individual with dyslexia whose variability was in this range. This pattern of results was replicated in Experiment 2 (Figs. S3-S4).

### Impaired online error correction underlies poor synchronization in ASD

Phase correction is the process of using the perceived error (deviation of the current tap from mean asynchrony) to adjust the timing of the next tap to be closer to the participant’s mean asynchrony. To test the efficiency of online phase correction we calculated the correlation between consecutive asynchronies. A positive correlation indicates under-correction, it means that deviations from mean asynchrony tend to persist across beats. Thus, weaker correction yields a larger correlation. All three groups showed a positive correlation (Fig. 2a-c, *r*_*CON*_ = 0.60, *r*_*DYS*_ = 0.59, *r*_*ASD*_ = 0.75), indicating that participants do not fully compensate for errors across consecutive beats (in line with Repp, 2011). Calculating single participant correlations, we found the largest correlation in the autism group (Fig. 2d, single participant correlations median [interquartile range]: neurotypical: 0.52 [0.23], dyslexia: 0.51 [0.27], autism: 0.69 [0.28], Kruskal Wallis test H(2)=8.86, p=0.012), indicating that they retain un-corrected errors longer than the other two groups. The difference between the groups was significant, and post-hoc comparisons showed that this is the result of a significant difference between the ASD group and both the neurotypical (p=0.033, Cliff’s delta = 0.35) and the dyslexia groups (p=0.017, Cliff’s delta = 0.39).

**Figure 2:**
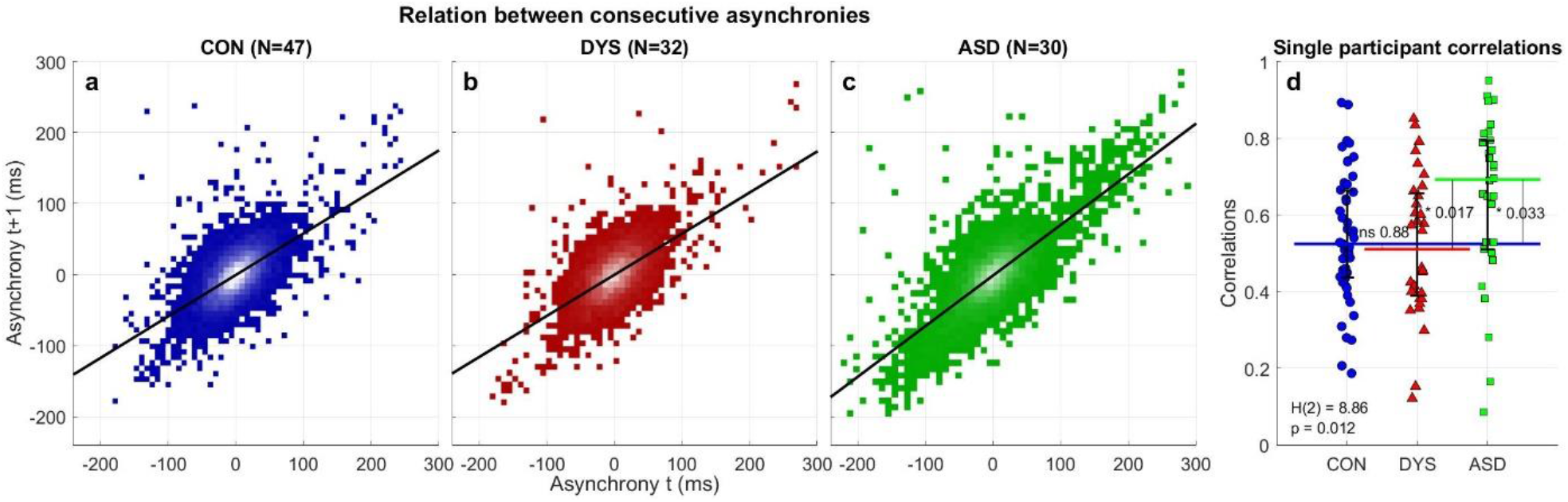
Correlation between consecutive asynchronies (errors) is highest in the ASD group revealing reduced online error correction. (a-c) Scatter plots showing correlations between consecutive asynchronies: (a) neurotypical, (b) dyslexia and (c) ASD. Individual asynchronies were plotted with respect to each participant’s mean asynchrony, yielding a mean of 0ms. Consecutive asynchronies are positively correlated in all groups. This positive correlation is largest in the ASD group, reflecting reduced online error correction. Luminance scale is equal in (a-c): white, the maximum count, is 165 in all graphs. (d) Single participant correlations also show the impairment in error correction for the ASD group compared with the neurotypical and dyslexia groups. The median of each group is denoted as a line of the same color; error bars denote interquartile range. Kruskal Wallis H-statistic and the corresponding p-value are plotted in the bottom-left corner; p-values of comparisons between groups are plotted next to the line connecting the groups’ medians.

To understand the dynamics of phase correction we used an autoregressive model to predict the current asynchrony with four predictors (the asynchronies at *t −* 1, *t −* 2, *t −* 3 and *t −* 4, see Methods). We administered the model for each participant separately and compared the fitted coefficients across the groups (Fig. S5). In accordance with the correlation results, we found a significant difference between the groups in the contribution of the most recent asynchrony to the current asynchrony (coefficient corresponding to the most recent trial - *b*_1_ mean ± SEM: neurotypical: 0.47 ± 0.03, dyslexia: 0.44 ± 0.02, autism: 0.59 ± 0.06; F(2,106)=4.083, p<0.02), indicating that the most recent error is a better predictor of current error in the ASD group compared with each of the other groups, i.e. it was corrected less. The contribution of earlier asynchronies decayed quickly in all groups, with only a small contribution for *b*_2_ (mean ± SEM: neurotypical: 0.08 ± 0.02, dyslexia: 0.08 ± 0.02, autism: 0.07 ± 0.04), whose magnitude did not differ between groups (p>0.4, uncorrected), and even smaller for *b*_3_ (mean ± SEM: neurotypical: 0.03 ± 0.01, dyslexia: 0.05 ± 0.02, autism: 0.02 ± 0.02). The contribution of *b*_4_ did not significantly differ from zero in any group (mean ± SEM: neurotypical: 0 ± 0.01, dyslexia: −0.01 ± 0.01, autism: −0.02 ± 0.02; Wilcoxon signed rank test, p>0.2 for all 3 groups). To summarize, error correction in paced tapping to an isochronous metronome is a fast mechanism which depends only on very recent history. This fast correction is reduced in ASD, and intact in dyslexia, in line with previous perceptual observations on the rate of perceptual update in both populations (Lieder et al., 2019).

### Modelling isochronous tapping reveals that only error (phase) correction is impaired in ASD

Impaired phase correction does not rule out that individuals with autism also have noisier representations of the metronome tempo (period), or “sloppier” production of motor commands (motor noise). To address this possibility, we used a well-established computational model of sensorimotor synchronization (Vorberg and Schulze 2002; Vorberg and Wing 1996; Jacoby et al., 2015). This model assumes that each tapping interval is the summation of three components: timekeeping of base tempo, time required for motor execution, and fraction of error (asynchrony) correction from the previous tap. Timekeeper is the process that maintains a representation of the external tempo (Ivry et al., 2002; Wing and Kristofferson 1973), and motor execution is the component that models the noise in executing motor commands. Previous work suggested that the motor noise, associated with each movement onset, and the timekeeper noise, associated with inter-beat intervals, can be distinguished from one another based on the covariance structure of the noise term (Wing & Kristofferson, 1973; Vorberg and Wing 1996; Vorberg and Schulze 2002). Formally, the model can be written as follows: *d*_*k*+1_ = (1 *− α*)*e*_*k*_ + *T*_*k*_ + *M*_*k*+1_ *− M*_*k*_ where *d*_*k*+1_ is the delay interval from the previous metronome stimulus (beat k) to the following finger tap (tap k+1), *e*_*k*_ is the asynchrony at beat k (see Fig. 1a), α denotes the proportion of correction of this asynchrony in tap k+1 (with a negative sign in the model since positive asynchrony deviations should be followed by shorter intervals), *T*_*k*_ is the participant’s current representation of the metronome tempo and *M*_*k*_ is the time of the motor response at time k (both including noise). Note that when *α* = 0 there is no correction and the previous asynchrony is carried to the next response. We fitted the model for each participant separately and compared the group parameters (Fig. 3).

**Figure 3:**
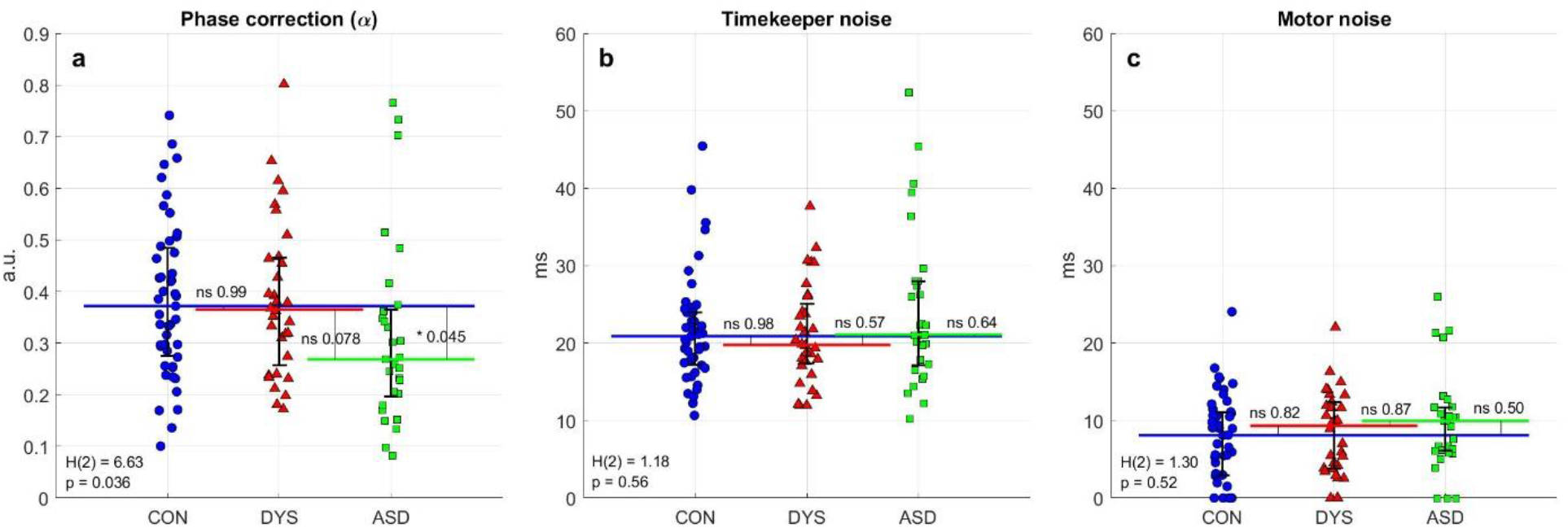
Parameters estimated for each participant, based on trial-by-trial modelling, show that individuals with autism have impaired error correction and intact timekeeper and motor noise. Results of the bGLS (bounded General Least Squares) estimation method (Jacoby et al., 2015) for a computational model of sensorimotor synchronization. (a) Error correction of phase difference – the fraction corrected (α) is significantly smaller in the ASD group. (b) Noise in keeping the metronome period, and (c) Motor noise do not differ between the groups. Each block was modeled separately, then parameters were averaged over the two assessment blocks. The median of each group is denoted as a line of the same color; error bars denote interquartile range. Kruskal Wallis H-statistic and corresponding p-value are in the bottom-left corner; p-values of comparisons between groups are next to the line connecting the groups’ medians.

Phase correction was (median [interquartile range]) 0.37 [0.21] in both the neurotypical and dyslexia groups, indicating that error was only partially corrected across consecutive taps, in line with the positive correlation we found (Fig. 2). Yet, phase correction was even smaller (0.27 [0.17]) in the autism group, with a significant group difference (Fig. 3a; Kruskal Wallis test H(2)=6.63, p=0.036). Post hoc analysis showed a significant difference between the neurotypical and autism groups (p=0.045, Cliff’s delta = 0.31) and a marginal difference between the dyslexia and autism groups (p=0.078, Cliff’s delta = 0.32), but no difference between the neurotypical and dyslexia groups (p>0.95). In contrast to phase correction, we found no group difference in the level of either timekeeping noise or motor noise (Fig 3b-c; time-keeper noise (median [interquartile range] (ms)): neurotypical: 20.9 [6.8], dyslexia: 19.8 [7.7], autism: 21.1 [10.8]; Kruskal Wallis test H(2)=1.18, p=0.56; motor noise (median [interquartile range] (ms)): neurotypical: 8.1 [8.2], dyslexia: 9.3 [8.6], autism: 10 [5.6]; Kruskal Wallis test H(2)=1.3, p=0.52). The specificity of the group difference to phase correction shows that the larger variability in the autism group does not stem from an elevated noise level in either motor or tempo keeping processes.

### Experiment 2 – tempo switches reveal impaired online updating to external changes in ASD

In the second finger-tapping experiment we asked whether individuals with autism or individuals with dyslexia have difficulties adapting to changing environments. We tested this by switching the tempo of the metronome, so that within each block the tempo alternated between two options (randomly every 8-12 intervals). We quantified the dynamics of updating to the new tempos in our three groups using both model-free and model-based analyses to uncover the underlying mechanisms.

#### Individuals with ASD fail to adapt to changes in the environment

Fig. 4 shows the timing of tapping in each population aligned to the onset of tempo change (left – acceleration, right – deceleration). For presentation purposes we aligned the pre-change delay interval with the metronome beat (cancelling the difference that originated from negative mean asynchrony, which varies across individuals). Since the time of tempo-change was not expected, the delay interval in the first beat after the change (beat 0) resembles that of the pre-change period. Following this initial surprise, participants updated their delay intervals to align with the new metronome tempo. This update was faster in the larger and more salient tempo changes (Repp, 2001; Repp & Keller, 2004): in the 90ms step-size (Fig. 4a-b), which is very salient, the neurotypical and dyslexia groups managed to synchronize to the new tempo after 1-2 metronome beats. This was not the case for the ASD group, which under-corrected in the first and second taps following the change and did not fully adapt even after 7 taps. Though this effect is clearest for the 90ms step-size, similar dynamics can be seen also in the 70ms step-size (Fig. 4c-d). The smaller, 50ms step-change (fig. 4e-f), was less salient and took marginally longer to adapt also for the dyslexia group compared with the neurotypical group, though the difference was not significant in any of our analyses (see following sections). Nonetheless, we cannot rule out the possibility that individuals with dyslexia manifest a small impairment in adjusting to small tempo switches.

**Figure 4:**
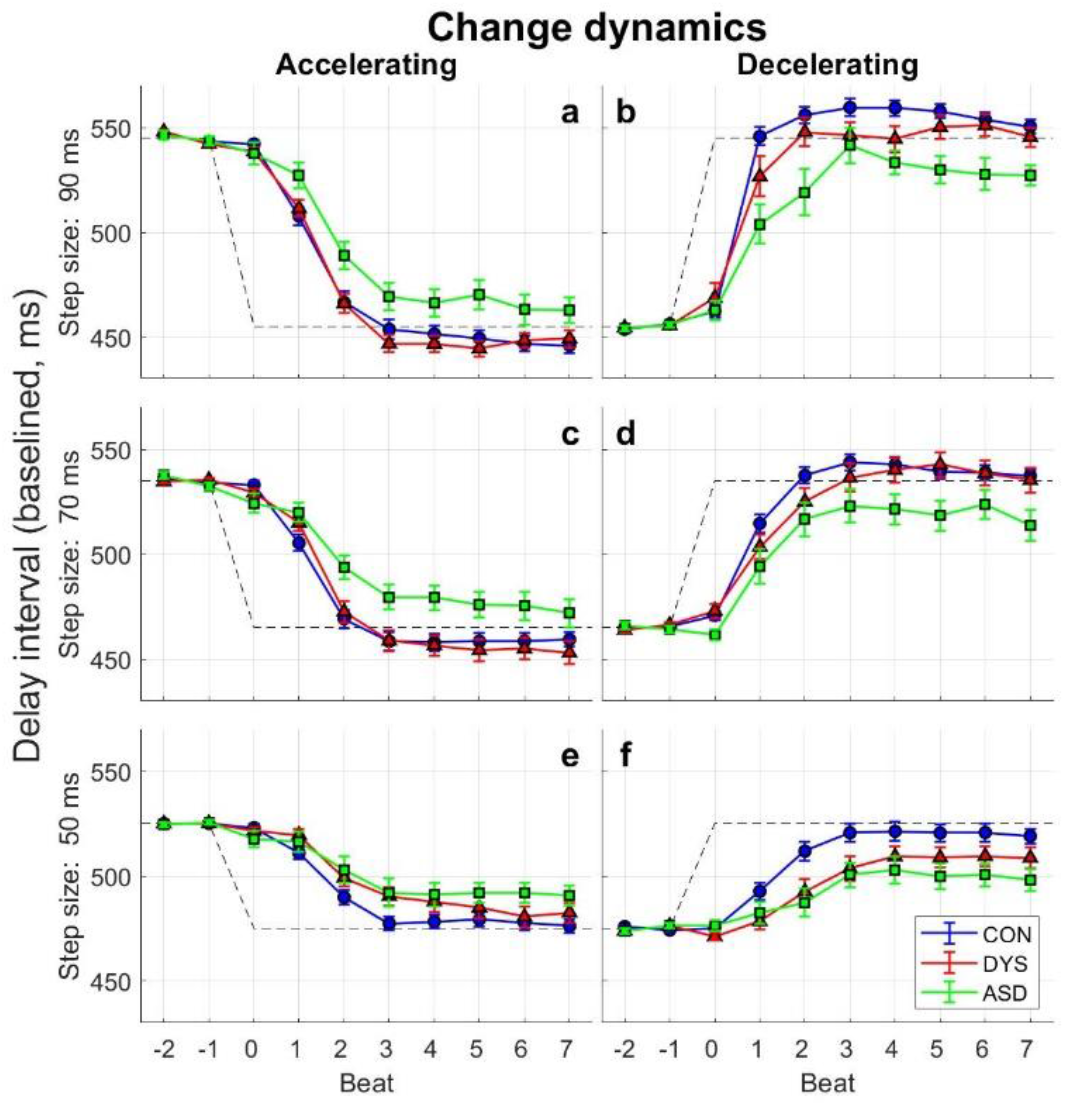
Individuals with autism adapt to changes in tempo only partially, even when changes are very salient. (a-b) 90ms step-size, (c-d) 70ms step-size and (e-f) 50ms step-size. In each panel the x-axis represents the metronome-beat number around the moment of tempo change (beat 0), and the y-axis measures the delay interval in each beat. Changes are quickly corrected, particularly for the larger (70, 90 ms) steps. Slower updates are seen for the smaller 50ms step changes, where neurotypicals take 3-4 steps to correct, and individuals with dyslexia take longer, perhaps since these steps are less salient. The difficulties of individuals with autism are seen in all step changes, their error is not fully corrected even within 7 taps. Delay interval is aligned to the pre-change metronome signal. Each participant tapped through 8-10 accelerations and decelerations in each condition. The values in the figure were calculated by first averaging responses within each participant and then across the group; error bars denote SEM across participants.

#### Individuals with ASD do not fully update to tempo changes even with longer time periods

To assess whether updating was attained several beats after the tempo-change, we calculated the distributions of the delay intervals in each of the metronome tempos, excluding the 4 beats immediately after the tempo change, where most tempo update takes place, as shown in Figure 4 (taking out 2-6 beats after the change produced similar statistics). If participants eventually adapt to the change in tempo, the two distributions should be highly separable. This was quantified using measurements from signal detection theory: sensitivity index (d’) and area under the curve (AUC) of the receiver-operator-characteristic (ROC). In the 90ms and 70ms step-sizes (Fig. 5a-h) we received comparable measurements for the neurotypical and dyslexia groups, and reduced values for the autism group, though in the 50ms step-size (Fig. 5i-l) the values of the dyslexia group are between those of the neurotypical and autism groups. This pattern was replicated when we looked at single participant values: there was a significant difference between the groups in all conditions (Kruskal Wallis Test; all p<0.01). Post-hoc comparisons showed a significant difference between the ASD and neurotypical groups (all p<0.01). The difference between the neurotypical and dyslexia groups was not significant in any step change (all p>0.4).

**Figure 5:**
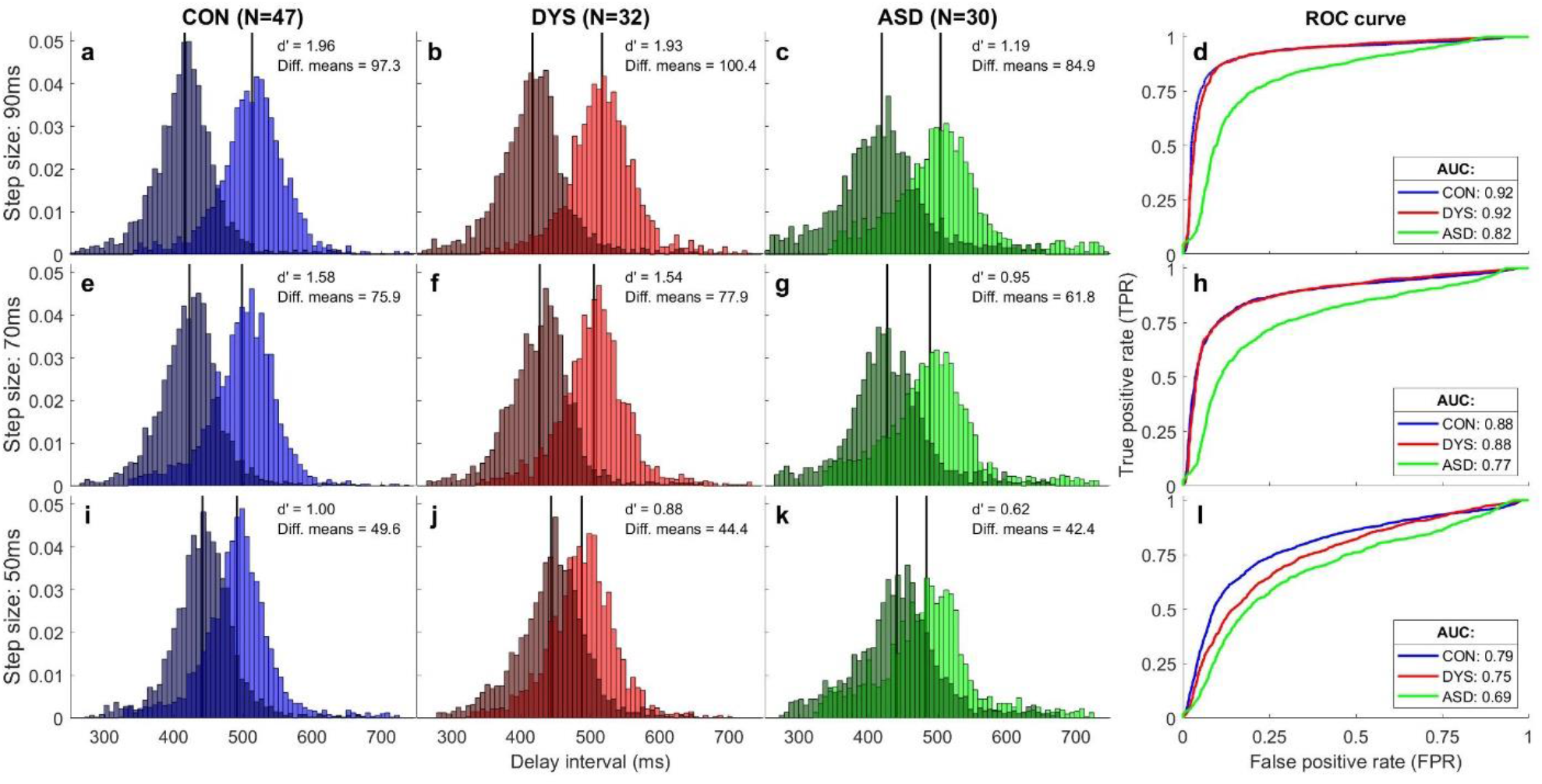
Distributions of delay intervals 5-12 taps after the tempo switch show that asynchronies in the autism group remain uncorrected. Panels (a-c), (e-g) and (i-k) show group delay intervals probability density functions separately for the longer tempo (light color) and shorter tempo (dark color), for each tempo-change condition (90ms – top, 70ms – middle, 50ms – bottom) and for each population (neurotypicals – blue, dyslexia – red, ASD – green). The mean of each distribution is denoted by a black vertical line. Values of d’ and the difference between the means are in the top right corner. Panels (d), (h) and (l) show for each group the receiver-operator-curves (ROC), and area under the curve (AUC) for classification between delay intervals under the two tempos. In the 70ms and 90ms step sizes, the measures of the dyslexia and neurotypical groups nearly overlap, while the values for the ASD group are smaller, reflecting reduced updating to changes in external tempo. For the small 50ms step-size the dyslexia group values are midway between the neurotypical and autism values, though this difference was not significant (see main text).

Importantly, d’ and AUC are both affected by the standard deviation (SD) of the distribution. Since the SD in ASD group is larger (Experiment 1), normalizing by SD would decrease d’ in this group more than in the other groups. In order to see if there is an impairment in the autism group on top of the increased variability, we used the difference between the means of the distributions without SD normalization. We found comparable values for the neurotypical and dyslexia groups, and smaller distances in the ASD group, for the 90ms and 70ms step-size conditions (Fig. 5a-h, Kruskal Wallis Test for single participants 90ms: p=0.007; 70ms: p=0.014). For the 50ms step-size (Fig. 5i-l) we found that the dyslexia group value is midway between that of the neurotypical and the ASD groups, as in other measures of small tempo changes (Kruskal Wallis Test p>0.2). Combined measures (formed by z-scoring each step-size condition, and then averaging over the different conditions) showed a significant difference between the groups in all measures (Kruskal-Wallis test all p<0.02), and post hoc comparisons showed no differences between the neurotypical and dyslexia groups (all p>0.4), but significant differences between the neurotypical and ASD groups (all p<0.02, all Cliff’s delta > 0.36) and between the dyslexia and ASD groups (p<0.05 for AUC and difference of means, p=0.08 for d’, all Cliff’s delta > 0.35). The retention of group difference across several taps partially stems from the over correction (for the large step-sizes) of the neurotypical and dyslexia groups (Figs 4 and 5).

#### Modelling the parameters underlying tempo switches reveals impaired period updating in ASD

To model the mechanisms underlying tempo changes we extended the computational model of Experiment 1. Following Schulze et al. 2005, we added a parameter that denotes the rate of the change in the internal representation of the metronome tempo as a fraction of the phase error that results from the tempo change. Formally, we add the following dynamics to the model: < *T*_*k*_ >= < *T*_*k−*1_ > *− βe*_*k*_, where *T*_*k*_ is the internal representation of the metronome tempo, *e*_*k*_ is the asynchrony at beat k, and β denotes the fraction of correction of the internal tempo, which is required due to the change in the metronome beat (period correction). Optimally < *T*_*k*_ > should be equal to the new metronome tempo, and this model estimates the rate of this process. The full model can be written as: *d*_*k*+1_ *− d*_*k*_ = *−βe*_*k*_ + (1 *− α*)(*e*_*k*_ *− e*_*k−*1_) + *Z*_*k*_ (Jacoby et al., 2015), where *d*_*k*+1_ is the participant’s delay interval between tap k+1 and stimulus k, *e*_*k*_ is the asynchrony at beat k, *Z*_*k*_ is a noise term incorporating motor and timekeeper noise and α and β are error correction terms (phase and period correction, respectively).

To enhance the model’s sensitivity to the changes, we used only the segments immediately before and after the tempo change. We fit the model to each tempo-change segment separately and averaged the resulting parameter values for each step-size. To obtain combined estimates we z-scored each parameter for each step-size condition (using the mean and standard deviation of all three groups combined) and averaged over the different conditions (Fig. 6). The ASD group had significantly smaller period correction (z-scored β, median [interquartile range]: neurotypical: 0.18 [1.1], dyslexia: −0.1 [1.06], autism: −0.35 [1.32]; Kruskal Wallis test H(2)=9.17, p=0.01). Post-hoc comparisons showed a significant difference between the ASD and neurotypical groups (p=0.007, Cliff’s delta = 0.4), with no other significant differences (both p>0.2). No differences were found in other estimated parameters (all p>0.5, see Fig. 6), including z-scored phase correction (α). To conclude, individuals with autism show an impairment in the initial updating of tempo, which is not fully corrected within the next 3-4 seconds (> 7 taps), as can be seen in Figure 4 and Figure 5.

**Figure 6:**
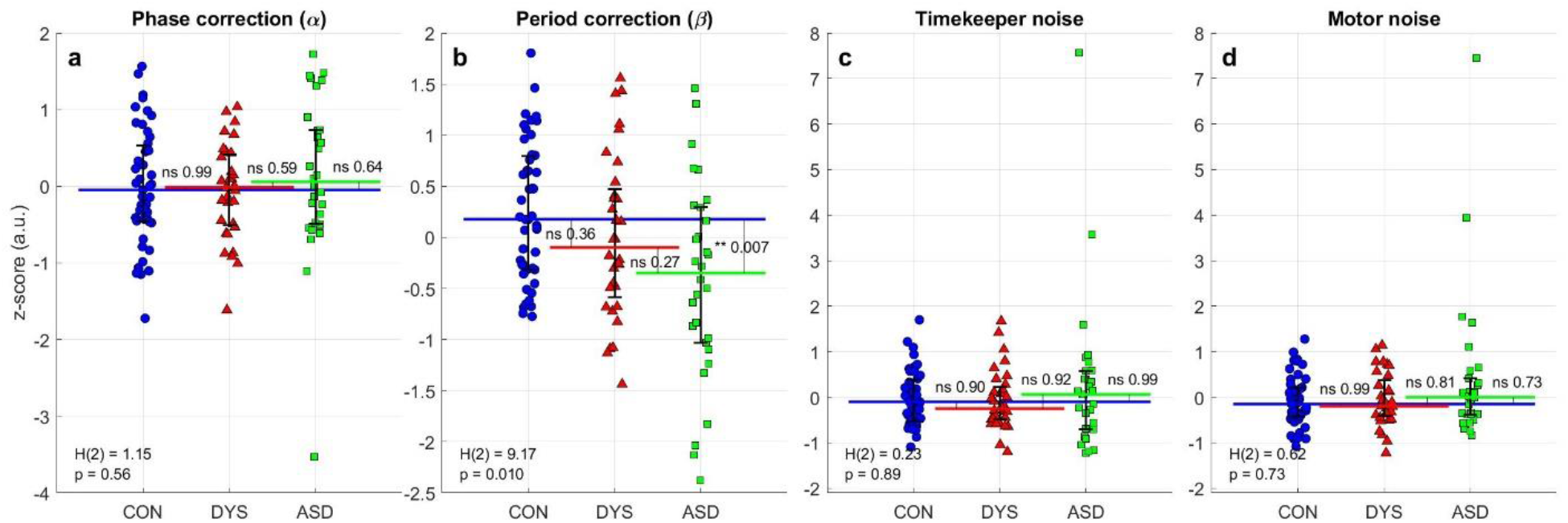
Trial-by-trial modelling of tempo changes shows that individuals with ASD have reduced period correction when faced with abrupt period changes. The panels show the results of the bGLS (bounded General Least Squares) estimation method (Jacoby et al., 2015) for the extended model of sensorimotor synchronization (Mates, 1994; Schulze et al., 2005). Each dot represents the combined value from all conditions of tempo-switches (50-90ms; after z-scoring using the mean and standard deviation of all groups). (a-b) Period correction (β) is decreased in the autism group compared with neurotypicals, while (α) is comparable across groups. (c) Timekeeper noise and (d) Motor noise estimates do not differ between the groups. The median of each group is denoted as a line of the same color; error bars denote interquartile range. Kruskal Wallis H-statistic and corresponding p-value are in the bottom-left corner; p-values of comparisons between groups are next to the line connecting the groups’ medians.

The extended model explained the data better than the model we used for the isochronous protocol, as evident by three different measures: Akaike information criterion (AIC is smaller than the original model for all subjects), Bayesian information criterion (BIC provided very strong evidence against the partial model for 108 of 109 subjects (Δ*BIC* > 10)) and likelihood ratio test (p<10^*−*10^ for all subjects). Moreover, our second correction term β, which denotes the update to the internal period, was significantly larger than the first correction term α in all groups and for all step-sizes (Wilcoxon signed-rank test, all p<0.015), indicating that our approach emphasizes correction of the internal tempo over standard phase correction.

Having found group differences in phase correction in a stationary environment (α, Experiment 1) and in period correction in the changing-tempo protocol (β, Experiment 2) we asked whether these two parameters denote separate mechanisms, or, alternatively, both reflect the same mechanism of online error correction. The relative contributions of the processes of correction for phase error and for period error are difficult to dissociate in a tempo-change paradigm, since these errors are temporally correlated (Repp, 2001; Jacoby and Repp, 2012). The large errors immediately following the tempo-change are always the summation of the error directly induced by the metronome’s tempo change (which requires a genuine period correction), and the error induced by the participant’s inability to predict the point of tempo-change (inducing an additional step-change phase error at beat zero). To resolve this ambiguity, we assessed the cross-participant correlation between the parameter of phase correction in Experiment 1 (Fig. 3a), and period correction in Experiment 2 (Fig. 6b). We found significant positive correlations in each of the three groups separately (Spearman correlations: *ρ*_*CON*_ = 0.47 (p<0.001), *ρ*_*DYS*_ = 0.42 (p<0.02) and *ρ*_*ASD*_ = 0.49 (p<0.008), Fig. 7) and when combining the groups (*ρ*_*ALL*_ = 0.51 (p<10^*−*7^)), with attenuated correlations all larger than 0.85, suggesting that almost all of the explained variability is shared between the two error correction terms. By contrast, there were no significant correlations between the other error correction parameters in all three groups (all *|ρ|* < 0.12, *p* > 0.4 for the correlation between the two error terms of Experiment 2, and the two estimations of phase correction). This combined pattern of correlations suggests that phase correction in Experiment 1 and period correction in Experiment 2 are manifestations of a common mechanism of online error correction, whose efficiency is reduced in autism, yielding slower correction rates in both fixed and changing environments.

**Figure 7:**
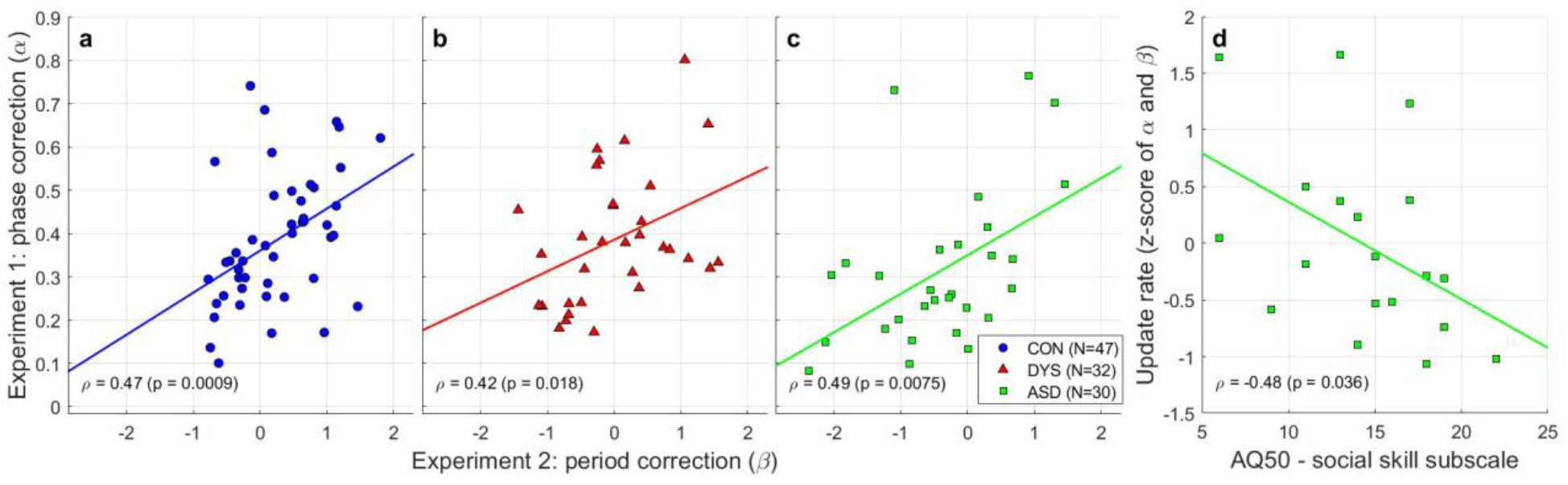
Rate of online error correction in stationary and in changing environments reflect a single underlying mechanism, which is slower in ASD and this mechanism is correlated with social skill. The estimated phase correction from Experiment 1, and the estimated period correction from Experiment 2 are highly correlated in all groups ((a) Neurotypical, (b) Dyslexia, (c) Autism), suggesting that both are manifestations of a common underlying mechanism of error correction which determines the speed of integrating new sensory data to guide behavior. The ASD group shows lower correction rates in both tasks. (d) The combined update rate (from both experiments) is correlated with the social-skill subscale of the AQ50 in the autism group. A higher AQ score indicates poorer skills. Correlation values are Spearman rank correlations, used due to the non-normality of the distributions of phase and period update values.

Finally, to test whether slower updating in autism is correlated with social difficulties, we administered the AQ50 (Autism Quotient) – a questionnaire based on self-report, aimed to assess the severity of difficulties (Baron-Cohen et al., 2001). It is composed of questions that aim to assess 5 separate subscales, which relate to different behavioral aspects that characterize autism. The most relevant subscale to our hypothesis is that assessing social skills. Nineteen participants with autism completed the questionnaire. As expected, their rate of update (the combined z-score of α in Experiment 1 and β in Experiment 2) and their social score (Fig. 7d, Spearman correlation *ρ* = *−*0.48, p=0.036) were significantly correlated.

## Discussion

We found that individuals with autism fail to synchronize their movements to external cues, whereas individuals with dyslexia synchronize adequately. Trial-by-trial computational modelling enabled us to precisely pinpoint the underlying deficit. We found that online error correction is impaired in the autism group while the level of noise in motor processing and internal timekeeping are adequate. The synchronization difficulties in a non-social context indicate that poor synchronization is not an outcome of a lack of social interest. These results extend the recently proposed “Slow-Updating” framework (Lieder et al., 2019), which asserts that individuals with autism are slow in integrating new sensory information, into the domain of sensorimotor synchronization, directly associated with social skills.

Importantly, our account of slower mechanisms of online error correction goes a long way towards explaining the muddle in the mixed literature assessing motor performance, sensorimotor performance, and even finger tapping in ASD. This literature characterized motor skills but did not study synchronization as the limiting bottleneck. For example, when individuals were asked to keep tapping after the metronome stops (unpaced tapping), performance of the ASD group was comparable to neurotypicals’ (Price et al., 2012; Ravizza et al., 2013). This seems surprising since these conditions are more cognitively demanding (Repp & Su, 2013). However, the bottleneck to performance here is keeping the previous tempo, i.e. robustness of working memory rather than synchronization with external stimuli. In such conditions we counter-intuitively predict that performance of individuals with ASD will be similar to that of neurotypical controls, since online error correction is not a limiting bottleneck. This is indeed the observation. Similarly, in demanding tasks that require more complicated learning mechanisms, and hence do not rely on online error correction, individuals with ASD are expected to show typical performance, which is indeed the case (Tryfon et al., 2017). However, when test conditions require online synchronization, their performance manifests elevated variability (Morimoto et al., 2018). Interestingly, in line with our analysis, error-related negativity (ERN) event potential has a lower amplitude and longer latency in ASD (Vlamings et al., 2008; Sokhadze et al., 2010). This ERP component is also associated with correction of large asynchronies in finger-tapping (Praamstra et al., 2003).

Our analyses suggest a novel mechanistic account to motor “clumsiness”, reported already in early descriptions of autism (Kanner, 1943; Asperger, 1944), and repetitively observed since then (Fournier et al., 2010; Gowen & Hamilton, 2013). We suggest that motor function is not inherently noisy in autism. Instead, the process of integrating sensory information into motor plans is slower in autism. Hence, while there is an essential sensory component to many movement forms (Gowen & Hamilton, 2013), we expect individuals with autism to experience the greatest difficulty when fast integration is required. This prediction is supported by recent reviews analyzing the core difficulties underlying poor sensorimotor integration in autism (Whyatt & Craig, 2013; Hannant et al., 2016a). Whyatt & Craig show that the motor deficit in autism is specific to tasks requiring fast sensorimotor integration, e.g., individuals with autism show a deficit in catching a ball, which requires rapid integration of visual information, while they show intact throwing, which is internally driven. Using a digitized version of the trace task from the M-ABC2, which requires participants to carefully navigate the cursor between the boundaries of a narrow line, they found that individuals with autism specifically failed in adjusting their behavior towards upcoming changes (corners). This specific difficulty explains their overall lower accuracy. Both reviews suggest that impaired sensorimotor synchronization may underlie all deficits found in autism spectrum disorder. We propose that sensorimotor synchronization is a specific manifestation of slow updating of internal models (Lieder et al., 2019).

This conclusion is further supported by seemingly unrelated findings in the visual domain. Individuals with autism show impaired processing of facial dynamics, but these impairments are alleviated when the rate of stimuli presentation is slowed down (Gepner et al., 2001; Gepner & Feron, 2009). Similarly, it has been shown that the difficulties of individuals with autism in performing tasks that require global motion perception are particularly severe when stimulus duration is short (Pellicano et al., 2005; Robertson et al., 2012; Robertson et al., 2014). This suggests that global perception per-se is not impaired but evolves more slowly (Robertson & Baron-Cohen, 2017). Another, seemingly unrelated, study used Bayesian modelling to understand the deficits of individuals with autism in a visual path integration task. Noel et al. (2019) found significantly higher variability in motor execution in the autism group compared to the neurotypical group, and their modelling framework revealed that individuals with autism are impaired in scaling their sensory likelihood function when executing the next action. Adequate scaling requires use of recent sensory information, particularly when the longer-term prior alone is inadequate to solve the specific task. Together, these studies support a multimodal account of slow updating in autism.

Within a Bayesian framework, our account proposes adequate use of slow accumulative information, yet poor use of recent cues. This account is in contrast to recently proposed Bayesian accounts which propose that individuals with autism overestimate the volatility of the environment (Palmer et al., 2017; Lawson et al., 2017), or that individuals with autism overweigh their prediction errors (Van de Cruys et al., 2014). According to these accounts, individuals with autism evaluate the environment’s statistics as changing more frequently than it actually does, and therefore they are expected to quickly update their internal model to meet their estimated degree of environmental change. We directly tested this prediction by using blocks with alternating tempos and found poor updating in the autism group. Therefore, while our results can be broadly viewed within a predictive coding context that emphasizes the role of sensory information and priors in guiding action (Clark, 2013), they do not support the increased volatility account or an account of overweighing prediction errors.

Our observation of no synchronization deficit in dyslexia are at odds with the temporal sampling framework of dyslexia (Goswami, 2011), which posits that individuals with dyslexia have problems with oscillatory entrainment, specifically in the delta range (1.5-4Hz). The theory predicts impairment in rhythmic motor performance at the tested range of 2Hz. Yet, early studies of individuals with dyslexia found no deficit in simple paced tapping tasks (Wolff et al., 1990; Tiffin-Richards et al., 2004). Follow-up studies (Thomson et al., 2006; Thomson & Goswami, 2008) had mixed results in paced finger tapping, and difficulties depended on the exact tempo around 2Hz. Still, we should note that we did find a subtle deficit in the dyslexia group in adapting to small tempo changes (50ms), though not in the isochronous condition. The difference from the neurotypical group was not significant in any of our analyses, but we cannot rule out a small deficit, since in this condition the dyslexia group’s performance did not significantly differ from that of the ASD group either.

The pattern of atypical performance of paced tapping among individuals with ASD suggests atypical development of one or two subcortical structures - the cerebellum and the basal ganglia. Both structures are related to temporal processing, though the precise role of each has been heavily disputed (Ivry et al., 2002; Diedrichsen et al., 2003; Repp, 2005; Repp & Su, 2013). Recent evidence suggests that the cerebellum is crucial for single-interval cueing tasks, namely, it keeps memory representation of intervals (‘timekeeping’), whereas the basal ganglia are more important for rhythmic tasks, namely, they play a role in online tracking (Teki et al., 2011; Breska & Ivry, 2018). According to this proposed division of labor, individuals with cerebellar lesions are expected to show increased noise levels due to impairment in representing the current metronome tempo, but intact phase or period correction (Ivry & Keele, 1989; Diedrichsen et al., 2003; though see Schwartze et al., 2016), while individuals with basal ganglia lesions are expected to show reduced error correction but intact noise levels (Diedrichsen et al., 2003; Schwartze et al., 2011). Our observation of intact timekeeper noise but reduced period and phase correction seems more consistent with a basal ganglia deficit. But, during development activity in many regions is likely to be influenced by any deficit, making it unlikely that a developmental atypicality will be consistently associated with one specific brain structure. Indeed, both the basal ganglia and the cerebellum display functional and anatomical abnormalities in autism spectrum disorders (Gowen & Miall, 2005; Fatemi et al., 2012; Subramanian et al., 2017).

To conclude, our results provide a novel computational account that bridges between previously reported atypical rate of perceptual updates, and social difficulties. It points to slow updating of motor plans as a core deficit in ASD, limiting sensorimotor synchronization and perhaps consequently impeding the development of social connectedness. Importantly, this characteristic is unique with respect to both neurotypical performance and to that of individuals with developmental dyslexia.

## Acknowledgements

We thank O. Guri and S. Granot for help collecting experimental data. We thank P. Dayan, U. Frith and Y. Hart for their support and helpful comments on the manuscript.

## Author contributions

NJ designed the experiments. TM initiated the collection of ASD data. OF, TM, TE and GV collected the data. Conceptualization: GV, NJ and MA. Formal analysis and methodology: GV and NJ. Writing – original draft: GV and MA. Writing – review & editing: GV, NJ and MA. Funding acquisition and supervision: MA.

## Data availability

All data and code are available from the authors upon request.

## Methods

### Participants

Neurotypical participants and participants with dyslexia were recruited through advertisements at the Hebrew University of Jerusalem and colleges near the university. Participants with ASD were recruited through clinics, designated facilities, and support groups. All participants were native Hebrew speakers (either born in Israel or immigrated to Israel before the age of 4 years). Performance on sensorimotor tasks is typically affected by musical background (Micheyl et al., 2006; Nahum et al., 2010; Repp 2005), and may affect different groups to a different extent (Oganian & Ahissar, 2012). We therefore recruited only participants with minimal musical experience (less than 2 years of self-reported musical education). All participants in the dyslexia group had been diagnosed by authorized clinicians as having a specific reading disability and all participants with ASD were diagnosed with autism spectrum disorders (including autism, asperger, and PDD-NOS) by expert clinicians. All participants completed a set of cognitive assessments, which evaluated general reasoning skills by the standard Block Design task (WAIS-IV) and reading abilities by pseudo-word and paragraph reading (details can be found in Lieder et al., 2019). They all performed the same protocol of finger tapping – Experiments 1 and 2.

Data were collected from 133 participants (56 neurotypical, 39 dyslexia and 38 autism). Of these, few were excluded, as follows. In order to match the groups on reasoning skills we excluded all participants with a Block Design score (Weschler, 2008) below 7 (1 dyslexia, 6 autism), and neurotypical and dyslexia participants with Block Design above 15 (7 neurotypical, 4 dyslexia). We assessed reading-related measures in the lab, leading us to exclude one neurotypical participant with exceptionally low pseudoword reading (41.7% accuracy, group average was 86.7%) and two participants with dyslexia with exceptionally high reading scores (one for 100% paragraph reading accuracy and one for 100% pseudoword reading accuracy). Finally, 3 participants were excluded due to extreme mean asynchrony values (>50ms, 1 neurotypical, 2 autism), suggesting a very different tapping strategy than that of other participants. The final group consisted of 109 participants (47 neurotypical, 30 females; 32 dyslexia). These groups were matched in age and reasoning skills, measured by the standard Block Design task. Results of these assessments are reported in Table S1.

All experiments were approved by the Ethics Committee of the Psychology Department of the Hebrew University and the Helsinki Ethics Committee of Sheba Hospital (required for testing individuals with ASD recruited through their adult clinic). All participants provided written informed consent and were financially compensated for their time and travel expenses.

### Finger Tapping Experimental Design

Participants heard a series of metronome beats and were asked to start tapping in synchrony with the metronome. To help participants synchronize, they were instructed to listen to the metronome first and tap after about 3 metronome beats (Repp, 2005). The metronome beats were heard through headphones at a comfortable presentation level. Tapping was performed on a custom-made wooden box, including a microphone which recorded the participant’s responses. We used either Focusrite Saffire 6 USB or Focusrite Scarlett 2i2 sound cards, which simultaneously recorded the output from the microphone installed inside the box and a split of the headphone signal, so that the playback latency and jitter could be estimated for each recording. Onsets were extracted from the stereo audio signal using a custom Matlab script. The overall latency and jitter obtained in this way, measured separately using calibration hardware, was about 1 millisecond (Elliott et al., 2018).

The task consisted of 12 blocks, each containing approximately 100 metronome beats. Rhythmic patterns consisted of short percussive sounds (“clicks”) lasting 55ms with an attack time of 5ms generated from amplitude modulated white noise. Blocks were separated by short pauses of 5 seconds. Participants had two breaks, after the 3^rd^ and 8^th^ blocks. Prior to the test procedure, all participants completed one block of practice. Blocks were separated into 6 conditions, and each was repeated twice. The first condition (Experiment 1) had an isochronous tempo of 2 Hz - beats were presented with an inter-onset-interval (IOI) of 500ms, known to be close to the optimal tempo for synchronization (Repp 2005; London 2002). The other five conditions (Experiment 2) were composed of alternating tempos. In each block the metronome tempo alternated between two options, which differed symmetrically from the baseline tempo of the isochronous condition (500ms): one tempo was faster than this baseline and the other was slower. Metronome changes occurred randomly every 8 to 12 intervals, thus both changes were repeated several times in each block. We used five different conditions with deviations ranging from ±5ms to ±45ms, in steps of 10ms: (1) 495ms and 505ms (±5ms, step size of 10ms), (2) 485ms and 515ms (±15ms, step size of 30ms), (3) 475ms and 525ms (±25ms, step size of 50ms), (4) 465ms and 535ms (±35ms, step size of 70ms) and (5) 455ms and 555ms (±55ms, step size of 90ms). Each block contained two types of changes: acceleration, when the tempo changed from the slow to the fast tempo, and deceleration, when it changed from the fast to the slow tempo. For example, in condition (3) the acceleration was a change from 525ms to 475ms and deceleration was the change from 475ms to 525ms. The 12 task blocks (including Experiment 1 and Experiment 2) were presented in one of four pseudorandomized orders.

As explained above, the tempo changes in Experiment 2 covered a broad range, and were chosen based on previous literature, which tested musicians or trained participants (e.g. Thaut et al., 1998). Our novice, musically untrained participants had markedly higher tracking thresholds – the two smaller step-changes were largely unnoticed by our participants (Figure S6). We therefore focused our analyses on the three larger step sizes shown in Figures 4-7. Importantly, the computational modelling results remain highly significant also when including the smaller tempo changes (Figure 6: β Kruskal Wallis test H(2)=10.41, p=0.005. Post hoc comparisons show a significant difference of autism and neurotypical groups (p=0.004); Figure 7: Spearman correlations *ρ*_*CON*_ = 0.55 (p<10^*−*4^), *ρ*_*DYS*_ = 0.4 (p=0.026) and *ρ*_*ASD*_ = 0.58 (p=0.0011)). For assessment of mean and standard deviation of the asynchrony we also analyzed the entire dataset (including the two smaller step sizes, Figures S2-S3).

### Finger Tapping Analyses

All analyses and statistical procedures were performed using Matlab. To measure synchronization, we used the time interval between the metronome stimulus and participant’s responses (termed asynchrony, Fig. 1a). Participants usually anticipate the metronome beat resulting in a negative mean asynchrony (Aschersleben & Prinz, 1995, Repp 2005). We excluded response taps that were outside a window of ±200 milliseconds surrounding metronome beats (Repp 2005). We denote by *S*_*t*_ and *R*_*t*_ the metronome onset and participant tap onset at beat number *t*, and denote by *e*_*t*_, *s*_*t*_, *r*_*t*_ and *d*_*t*_ the asynchrony (error), inter-beat (stimulus) interval, inter-tap (response) interval and delay interval, respectively (Fig. 1a). Formally: (1) *e*_*t*_ = *R*_*t*_ *− S*_*t*_ (2) *s*_*t*_ = *S*_*t*_ *− S*_*t−*1_ (3) *r*_*t*_ = *R*_*t*_ *− R*_*t−*1_ (4) *d*_*t*_ = *R*_*t*_ *− S*_*t−*1_. In the second experiment, perturbations of the metronome tempo occurred at unexpected time points, therefore we computed an adjusted asynchrony (*e*′_*t*_), which is the asynchrony after removing the contribution of the unexpected perturbation: *e*′_*t*_ = *e*_*t*_ + *s*_*t*_ *− s*_*t−*1_ (Figures S1-S2).

We computed:

1. Mean asynchrony = < *e*′ >
2. Standard deviation of the asynchrony = *std*(*e*′)

Results (Fig. 1b-c) were averaged over the two repetitions of each condition.

#### Experiment 1

##### Autocorrelation analysis

as a first approach to assess rate of error correction we computed the correlations between consecutive asynchronies (*e*_*t*_). These were calculated both at the population level (pooling over participants), and for each participant separately. In both cases, the mean for each block was subtracted from the values of that block, before the participant values were pooled together, or the two repetitions of the isochronous task pooled together. We used the Pearson correlation coefficient between *e*_*t*_*−*< *e* > and *e*_*t−*1_*−*< *e* >, where < *e* > is calculated per block.

##### Auto-regressive model

to study the timescale of serial dependence in tapping tasks, we used an autoregressive model with 4 predictors. Before estimating the coefficients, we subtracted the mean asynchrony from each block and then pooled both blocks together, as in the auto-correlations analysis. We then regressed the asynchrony at time t on the 4 preceding asynchronies:

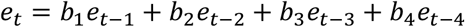

We stopped with 4 predictors since this was the first component where the dependence in all 3 groups was not significantly different from 0 (Wilcoxon signed rank test, p>0.2 for all 3 groups).

##### Computational model of sensorimotor synchronization

to test whether individuals with autism show noisier representations or “sloppier” motor production we used a computational model of sensorimotor synchronization (Vorberg and Wing 1996; Vorberg and Schulze 2002; Wing and Kristofferson 1973) where each interval is influenced by phase correction, timekeeping and motor execution. Phase correction is modeled as the proportion of the previous asynchrony error that is corrected in the execution of the following interval. Timekeeping is a process that maintains a representation of the tempo (Ivry et al., 2002), and the motor execution term models the execution noise. Formally, the model can be written as follows:

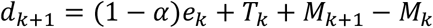

Where *d*_*k*+1_ is the participant’s delay interval between taps *k+1* and stimulus *k, e*_*k*_ is the asynchrony at beat *k, T*_*k*_ is the participant’s representation of the metronome tempo (timekeeper) and *M*_*k*_ is the motor execution (including noise) at time k. α denotes the degree of phase correction (with a negative sign in the model since positive asynchrony deviations should be followed by shorter intervals). The timekeeper (*T*_*k*_) is modeled as a Gaussian centered around the tempo of the metronome, and the motor execution component is modelled as a Gaussian around zero. The model takes as an input the empirical *d*_*k*_ and *e*_*k*_ and estimates the timekeeper and motor components (*T*_*k*_ and *M*_*k*_) and phase correction (α). Motor and timekeeping noise are defined as *std*(*M*_*k*_) and *std*(*T*_*k*_) respectively. As demonstrated in Jacoby et al. 2015, under the assumption of an upper bound on the magnitude of the motor noise (*std*(*M*_*k*_) < *std*(*T*_*k*_)), the parameters of the model can be reliably estimated, since each parameter has a unique contribution to the auto-covariance function of the signal (Jacoby et al., 2015). We fit the model for each block separately and averaged the two repetitions of the isochronous condition (Fig. 3). Blocks with more than 40% missing values (either due to participants missing a beat or tapping outside the ±200ms window from the metronome beat) were excluded from this analysis. Importantly, the Jacoby et al., 2015 version of the algorithm for parameter extraction does not enable fitting with missing values. We adapted the algorithm to enable fitting the model with missing data (supplementary material).

#### Experiment 2

##### Response dynamics to changes in tempo

to assess how participants respond to changes in tempo we aligned the participants’ responses to the tempo change and averaged each participant’s responses to acceleration and deceleration separately (Fig. 4). For presentation purposes we aligned the baseline delay intervals to the metronome tempo by subtracting the average asynchrony in the two intervals before the change from the entire segment. We included only transitions where all responses were available from two taps before the change (to establish a baseline asynchrony) to seven taps after the change (to assess the full progression of the adaptation procedure). Transitions with missing values in this range, or that were too close to the start or end of the block were excluded. Fig. 4 shows only participants with at least 2 repetitions of a given transition magnitude and direction.

##### Update to changes after several taps

We used the distributions of the delay intervals under each metronome tempo separately (pooling over the two repetitions of each condition). We excluded the 4 beats immediately following the change (including the moment of change; taking out 2-6 beats after the change produced similar statistics). If participants fully adapt to the change, the two distributions should be highly separable. We quantified this using 3 measures:

1. Sensitivity index, or d’: the distance between the means normalized by the standard deviations:

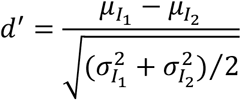

Where 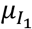 and Where 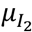 are the means of distributions 1 and 2, and 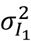 and 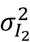 are the variances.
2. AUC (area under the curve): we create a receiver operating characteristic (ROC) curve by varying the threshold of a binary classifier designed to discriminate between the two distributions (so a delay interval below the threshold is marked as short tempo, and a delay interval above the threshold is marked as long tempo). For each threshold we calculate the percentage of true positives (TPR = true positive rate, delay intervals in the short tempo that were classified correctly) and false positives (FPR = false positive rate, delay intervals in the long tempo that were classified incorrectly as short tempo). AUC is the area under the curve of TPR vs FPR:

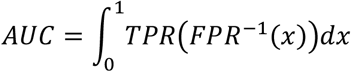
3. Difference between the means of the distributions, without normalizing it by the standard deviations. Meaning:

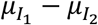

Where 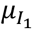 and 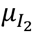 are the means of distributions 1 and 2.

##### Combined measures

In all cases combined measures were calculated by z-scoring each step-size separately, then averaging over the different step-sizes. Z-scoring was performed using the mean and standard deviation of the 3 groups together.

##### Extended computational model of sensorimotor synchronization

To understand whether individuals with autism manifest an impairment in their response to external changes, we extended the computational model of Experiment 1 (Schulze et al. 2005; Mates 1994), since the original model cannot capture responses to tempo-changes (the timekeeper distribution is set to a fixed mean). The extended model incorporates all parameters of the isochronous model (phase correction, timekeeper noise and motor noise) and an additional parameter which denotes the rate of change in the internal representation of the metronome tempo as a fraction of the resulting phase error. In this model the timekeeper is updated using the following dynamics:

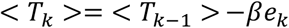

Where β denotes the fraction of correction of the internal tempo for a given tap (period correction), and < *T*_*k*_ > is the representation of the metronome tempo. The full model can be written as: *d*_*k*+1_ *− d*_*k*_ = *−βe*_*k*_ + (1 *− α*)(*e*_*k*_ *− e*_*k−*1_) + *Z*_*k*_ (Jacoby et al., 2015), where α and β are the phase and period correction respectively, *d*_*k*+1_ is the participant’s delay interval between tap k+1 and metronome beat k, *e*_*k*_ is the asynchrony at beat k and *Z*_*k*_ = *T*_*k*_ *− T*_*k−*1_ + *M*_*k*+1_ *−* 2*M*_*k*_ + *M*_*k−*1_ is a noise term incorporating motor and timekeeper noise (defined in the same way as in the partial model). The link to the model from Experiment 1 becomes clear if we take the difference between the model equation at time k+1 and at time k: *d*_*k*+1_ *− d*_*k*_ = (1 *− α*)*e*_*k*_ *−* (1 *− α*)*e*_*k−*1_ + *Z*_*k*_ = (1 *− α*)(*e*_*k*_ *− e*_*k−*1_) + *Z*_*k*_. Namely, we receive the same model as the model for Experiment 2, with the parameter β set to 0.

We fit the model separately for each tempo-change segment (from two beats before the change to seven beats following the change, see section *Response dynamics to changes in tempo*) and average the resulting model estimates. The fit was performed using the version of the model described in the appendix of Jacoby et al., 2015. We excluded segments with missing values, as described in *Response dynamics to changes in tempo*. For each step-size we excluded participants with less than 3 complete change segments (in both acceleration and deceleration).

##### Model comparison

The extended computational model was compared to the original partial model from experiment 1 using Akaike Information Criterion (AIC), Bayesian Information Criterion (BIC) and likelihood ratio test. AIC was computed according to the following formula:

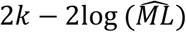

BIC was computed according to the following formula:

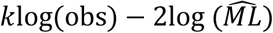

Where k is the number of model parameters, obs is the number of observations, and 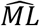 is the maximum likelihood estimate of the model, calculated by treating the model as a multivariate Gaussian with the estimated covariance matrix (Jacoby et al., 2015). For both AIC and BIC the log likelihood was summed across segments and between step-sizes (since the segments are independent). According to Kass & Raftery (1993) Δ*BIC* > 10 is considered very strong evidence against the model with the higher BIC.

## References

Ahissar, Merav, et al. “Auditory processing parallels reading abilities in adults.” Proceedings of the National Academy of Sciences 97.12 (2000): 6832–6837.

American Psychiatric Association. (2013). Diagnostic and statistical manual of mental disorders (5th ed.). Arlington, VA: American Psychiatric Publishing

Aschersleben, Gisa, and Wolfgang Prinz. “Synchronizing actions with events: The role of sensory information.” Perception & Psychophysics 57.3 (1995): 305–317.

Asperger, H. (1944). Autistic psychopathy in children. Translated in U. Frith (1991) Autism and Asperger’s syndrome.

Baron-Cohen, Simon, et al. “The autism-spectrum quotient (AQ): Evidence from asperger syndrome/high-functioning autism, malesand females, scientists and mathematicians.” Journal of autism and developmental disorders 31.1 (2001): 5–17.

Breska, Assaf, and Richard B. Ivry. “Double dissociation of single-interval and rhythmic temporal prediction in cerebellar degeneration and Parkinson’s disease.” Proceedings of the National Academy of Sciences 115.48 (2018): 12283–12288.

Chevallier, Coralie, et al. “The social motivation theory of autism.” Trends in cognitive sciences 16.4 (2012): 231–239.

Cirelli, Laura K. “How interpersonal synchrony facilitates early prosocial behavior.” Current opinion in psychology 20 (2018): 35–39.

Clark, Andy. “Whatever next? Predictive brains, situated agents, and the future of cognitive science.” Behavioral and brain sciences 36.3 (2013): 181–204.

Diedrichsen, Jörn, Richard B. Ivry, and Jeff Pressing. “Cerebellar and basal ganglia contributions to interval timing.” Functional and neural mechanisms of interval timing (2003): 457–481.

Fatemi, S. H., Aldinger, K. A., Ashwood, P., Bauman, M. L., Blaha, C. D., Blatt, G. J., … & Estes, A. M. (2012). Consensus paper: pathological role of the cerebellum in autism. The Cerebellum, 11(3), 777–807.

Fournier, Kimberly A., et al. “Motor coordination in autism spectrum disorders: a synthesis and meta-analysis.” Journal of autism and developmental disorders 40.10 (2010): 1227–1240.

Gepner, Bruno, Christine Deruelle, and Stanislas Grynfeltt. “Motion and emotion: A novel approach to the study of face processing by young autistic children.” Journal of autism and developmental disorders 31.1 (2001): 37–45.

Gepner, Bruno, and François Féron. “Autism: a world changing too fast for a mis-wired brain?.” Neuroscience & Biobehavioral Reviews 33.8 (2009): 1227–1242.

Germanò, Eva, Antonella Gagliano, and Paolo Curatolo. “Comorbidity of ADHD and dyslexia.” Developmental neuropsychology 35.5 (2010): 475–493.

Goswami, U. (2011). A temporal sampling framework for developmental dyslexia. Trends in cognitive sciences, 15(1), 3–10.

Goswami, U. (2015). Sensory theories of developmental dyslexia: three challenges for research. Nature Reviews Neuroscience, 16(1), 43.

Gowen, Emma, and R. Chris Miall. “Behavioural aspects of cerebellar function in adults with Asperger syndrome.” he Cerebellum 4.4 (2005): 279–289.

Gowen, Emma, and Antonia Hamilton. “Motor abilities in autism: a review using a computational context.” Journal of autism and developmental disorders 43.2 (2013): 323–344.

Hannant, P., Tavassoli, T., & Cassidy, S. (2016a). he role of sensorimotor difficulties in autism spectrum conditions. Frontiers in neurology, 7, 124.

Hannant, Penelope, et al. “Sensorimotor difficulties are associated with the severity of autism spectrum conditions.” Frontiers in integrative neuroscience 10 (2016b): 28.

Hove, Michael J., and Jane L. Risen. “It’s all in the timing: Interpersonal synchrony increases affiliation.” Social cognition 27.6 (2009): 949–960.

Ivry, R. B., & Keele, S. W. (1989). Timing functions of the cerebellum. Journal of cognitive neuroscience, 1(2), 136–152.

Ivry, Richard B., et al. “The cerebellum and event timing.” Annals of the new York Academy of Sciences 978.1 (2002): 302–317.

Jacoby, Nori, and Bruno H. Repp. “A general linear framework for the comparison and evaluation of models of sensorimotor synchronization.” Biological cybernetics 106.3 (2012): 135–154.

Jacoby, Nori, et al. “Parameter estimation of linear sensorimotor synchronization models: phase correction, period correction, and ensemble synchronization.” Timing & Time Perception 3.1-2 (2015): 52–87.

Jaswal, Vikram K., and Nameera Akhtar. “Being versus appearing socially uninterested: Challenging assumptions about social motivation in autism.” Behavioral and Brain Sciences 42 (2019).

Kanner, Leo. “Autistic disturbances of affective contact.” Nervous child 2.3 (1943): 217–250.

Kokal, Idil, et al. “Synchronized drumming enhances activity in the caudate and facilitates prosocial commitment-if the rhythm comes easily.” PLoS One 6.11 (2011): e27272.

Lawson, Rebecca P., Christoph Mathys, and Geraint Rees. “Adults with autism overestimate the volatility of the sensory environment.” Nature neuroscience 20.9 (2017): 1293.

Lieder, Itay, et al. “Perceptual bias reveals slow-updating in autism and fast-forgetting in dyslexia.” Nature neuroscience 22.2 (2019): 256.

Mates, J. (1994). A model of synchronization of motor acts to a stimulus sequence. Biological cybernetics, 70(5), 463–473.

Morimoto, Chie, et al. “Temporal processing instability with millisecond accuracy is a cardinal feature of sensorimotor impairments in autism spectrum disorder: analysis using the synchronized finger-tapping task.” Journal of autism and developmental disorders 48.2 (2018): 351–360.

Noel, Jean-Paul, et al. “Increased variability but intact integration during visual navigation in Autism Spectrum Disorder.” bioRxiv (2019).

Novembre, Giacomo, Zoe Mitsopoulos, and Peter E. Keller. “Empathic perspective taking promotes interpersonal coordination through music.” Scientific reports 9.1 (2019): 1–12.

Palmer, Colin J., Rebecca P. Lawson, and Jakob Hohwy. “Bayesian approaches to autism: Towards volatility, action, and behavior.” Psychological bulletin 143.5 (2017): 521.

Pellicano, Elizabeth, et al. “Abnormal global processing along the dorsal visual pathway in autism: a possible mechanism for weak visuospatial coherence?.” Neuropsychologia 43.7 (2005): 1044–1053.

Praamstra, P., et al. “Neurophysiological correlates of error correction in sensorimotor-synchronization.” Neuroimage 20.2 (2003): 1283–1297.

Price, Kelly J., Dorothy Edgell, and Kimberly A. Kerns. “Timing deficits are implicated in motor dysfunction in Asperger’s Syndrome.” Research in Autism Spectrum Disorders 6.2 (2012): 857–860.

Ravizza, Susan M., et al. “Restricted and repetitive behaviors in autism spectrum disorders: The relationship of attention and motor deficits.” Development and psychopathology 25.3 (2013): 773–784.

Repp, Bruno H. “Processes underlying adaptation to tempo changes in sensorimotor synchronization.” Human movement science 20.3 (2001): 277–312.

Repp, Bruno H., and Peter E. Keller. “Adaptation to tempo changes in sensorimotor synchronization: Effects of intention, attention, and awareness.” Quarterly Journal of Experimental Psychology Section A 57.3 (2004): 499–521.

Repp, Bruno H. “Sensorimotor synchronization: a review of the tapping literature.” Psychonomic bulletin & review 12.6 (2005): 969–992.

Repp, B. H. (2011). Comfortable synchronization of drawing movements with a metronome. Human Movement Science, 30, 18–39

Repp, Bruno H., and Yi-Huang Su. “Sensorimotor synchronization: a review of recent research (2006–2012).” Psychonomic bulletin & review 20.3 (2013): 403–452.

Robertson, Caroline E., et al. “Atypical integration of motion signals in autism spectrum conditions.” PloS one 7.11 (2012): e48173.

Robertson, Caroline E., et al. “Global motion perception deficits in autism are reflected as early as primary visual cortex.” Brain 137.9 (2014): 2588–2599.

Robertson, Caroline E., and Simon Baron-Cohen. “Sensory perception in autism.” Nature Reviews Neuroscience 18.11 (2017): 671–684.

Rogers, S. J., & Ozonoff, S. (2005). Annotation: What do we know about sensory dysfunction in autism? A critical review of the empirical evidence. Journal of Child Psychology and Psychiatry, 46(12), 1255–1268.

Schulze, H.-H., Cordes, A., & Vorberg, D. (2005). Keeping synchrony while tempo changes: Accelerando and ritardando. Music Perception, 22, 461–477.

Schwartze, Michael, et al. “The impact of basal ganglia lesions on sensorimotor synchronization, spontaneous motor tempo, and the detection of tempo changes.” Behavioural brain research 216.2 (2011): 685–691.

Schwartze, Michael, Peter E. Keller, and Sonja A. Kotz. “Spontaneous, synchronized, and corrective timing behavior in cerebellar lesion patients.” Behavioural brain research 312 (2016): 285–293.

Smith, Kimberly RM, and Johnny L. Matson. “Psychopathology: Differences among adults with intellectually disabled, comorbid autism spectrum disorders and epilepsy.” Research in Developmental Disabilities 31.3 (2010): 743–749.

Sokhadze, Estate, et al. “Impaired error monitoring and correction function in autism.” Journal of Neurotherapy 14.2 (2010): 79–95.

Subramanian, Krishna, et al. “Basal ganglia and autism–a translational perspective.” Autism Research 10.11 (2017): 1751–1775.

Tallal, Paula. “Improving language and literacy is a matter of time.” Nature Reviews Neuroscience 5.9 (2004): 721.

Tarr, Bronwyn, Jacques Launay, and Robin IM Dunbar. “Music and social bonding:”self-other” merging and neurohormonal mechanisms.” Frontiers in psychology 5 (2014): 1096.

Teki, Sundeep, et al. “Distinct neural substrates of duration-based and beat-based auditory timing.” Journal of Neuroscience 31.10 (2011): 3805–3812.

Thomson, Jennifer M., et al. “Auditory and motor rhythm awareness in adults with dyslexia.” Journal of research in reading 29.3 (2006): 334–348.

Thomson, Jennifer M., and Usha Goswami. “Rhythmic processing in children with developmental dyslexia: auditory and motor rhythms link to reading and spelling.” Journal of Physiology-Paris 102.1-3 (2008): 120–129.

C. Tiffin-Richards, Margaret, et al. “Time reproduction in finger tapping tasks by children with attention-deficit hyperactivity disorder and/or dyslexia.” Dyslexia 10.4 (2004): 299–315.

Tryfon, Ana, et al. “Auditory-motor rhythm synchronization in children with autism spectrum disorder.” Research in Autism Spectrum Disorders 35 (2017): 51–61.

Valdesolo, Piercarlo, and David DeSteno. “Synchrony and the social tuning of compassion.” Emotion 11.2 (2011): 262.

Van de Cruys, Sander, et al. “Precise minds in uncertain worlds: Predictive coding in autism.” Psychological review 121.4 (2014): 649.

Vlamings, Petra HJM, et al. “Reduced error monitoring in children with autism spectrum disorder: an ERP study.” European Journal of Neuroscience 28.2 (2008): 399–406.

Vorberg, Dirk, and Alan Wing. “Modeling variability and dependence in timing.” Handbook of perception and action. Vol. 2. Academic Press, 1996. 181–262.

Vorberg, Dirk, and Hans-Henning Schulze. “Linear phase-correction in synchronization: Predictions, parameter estimation, and simulations.” Journal of Mathematical Psychology 46.1 (2002): 56–87.

Wing, Alan M., and Alfred B. Kristofferson. “Response delays and the timing of discrete motor responses.” Perception & Psychophysics 14.1 (1973): 5–12.

Whyatt, C., & Craig, C. (2013). Sensory-motor problems in Autism. Frontiers in integrative neuroscience, 7, 51.

Wolff, Peter H., et al. “Rate and timing precision of motor coordination in developmental dyslexia.” Developmental Psychology 26.3 (1990): 349.

## Additional references for the methods section

Elliott, Mark T., et al. “Analysing Multi-person Timing in Music and Movement: Event Based Methods.” Timing and Time Perception: Procedures, Measures, & Applications. BRILL, 2018. 177–215.

Kass, Robert E., and Adrian E. Raftery. “Bayes factors.” Journal of the american statistical association 90.430 (1995): 773–795.

London, Justin. “Cognitive constraints on metric systems: Some observations and hypotheses.” Music Perception: An Interdisciplinary Journal 19.4 (2002): 529–550.

Micheyl, Christophe, et al. “Influence of musical and psychoacoustical training on pitch discrimination.” Hearing research 219.1-2 (2006): 36–47.

Nahum, Mor, et al. “From comparison to classification: A cortical tool for boosting perception.” Journal of Neuroscience 30.3 (2010): 1128–1136.

Oganian, Yulia, and Merav Ahissar. “Poor anchoring limits dyslexics’ perceptual, memory, and reading skills.” Neuropsychologia 50.8 (2012): 1895–1905.

Thaut, M. H., Miller, R. A., & Schauer, L. M. (1998). Multiple synchronization strategies in rhythmic sensorimotor tasks: phase vs period correction. Biological cybernetics, 79(3), 241–250.

Weschler, D. Wechsler Adult Intelligence Scale. 4th edn (Pearson: London, 2008).

